# When the liver is in poor condition, so is the heart - cardiac remodelling in MASH mouse models

**DOI:** 10.1101/2024.05.01.592038

**Authors:** Sebastian Bott, Justine Lallement, Alice Marino, Evangelos P. Daskalopoulos, Christophe Beauloye, Hrag Esfahani, Chantal Dessy, Isabelle Anne Leclercq

**Affiliations:** Laboratory of Hepato-Gastroenterology, Institut de Recherche Experimentale et Clinique, Universite catholique de Louvain, Brussels; Pole of Pharmacology and Therapeutics, Institut de Recherche Experimentale et Clinique, Universite catholique de Louvain, Brussels, Belgium; Pole of Cardiovascular Research, Institut de Recherche Experimentale et Clinique, Universite catholique de Louvain, Brussels, Belgium; Division of Cardiology, Cliniques Universitaires Saint-Luc, Brussels, Belgium; Platform of Integrated Physiology, Institut de Recherche Experimentale et Clinique, Universite catholique de Louvain, Brussels, Belgium

**Author notes:** **Corresponding authors,** Chantal Dessy, FATH, Avenue Hippocrate 57/B1.57.04, 1200 Woluwe-Saint-Lambert, Brussels, Isabelle A. Leclercq, GAEN, Avenue Mounier 53/B1.52.01, 1200 Woluwe-Saint-Lambert, Brussels.

**Keywords:** MASLD, MASH, CVD, cardiac hypertrophy, adverse cardiac remodelling, foetal gene reprogramming, *foz/foz*, C57BL/6J

## Abstract

Metabolic dysfunction-associated steatohepatitis (MASH) confers a risk for cardiovascular diseases in patients. Animal models may help exploring the mechanisms linking liver and heart diseases. Hence, we explored the cardiac phenotype in MASH mouse models.

**Methods:** We evaluated pathological alterations in liver and heart in *foz/foz* mice fed a high fat diet for 24 or 60 weeks and in C57BL/6J mice fed a high fat, high fructose diet for 60 weeks. Angiotensin ll (Angll) was used as an additional cardiovascular stressor.

**Results:** F*oz/foz* mice with fibrosing MASH developed cardiac hypertrophy with adverse cardiac remodelling not seen in WT similarly fed the high fat diet. Angll caused hypertension and upregulated the expression of genes contributing to pathological cardiac hypertrophy (*Nppa*, *Myh7*) more severely so in *foz/foz* mice than in controls. After 60 weeks of HFD, while liver disease had progressed to burn-out non steatotic MASH with hepatocellular carcinoma in 50% of the animals, the cardiomyopathy did not. In an independent model (C57BL/6J mice fed a high fat, high fructose diet), moderate fibrosing MASH is associated with cardiac fibrosis and dysregulation of genes involved in pathological remodelling (*Collal*, *Col3al*, *Vim*, *Myh6*, *Slc2al*).

**Conclusion:** Animals with MASH present consistent adverse structural changes in the heart with no patent alteration of cardiac function even when stressed with exogenous Angll. Liver disease, and not overfeeding or aging alone, is associated with this cardiac phenotype. Our findings confirm *foz/foz* mice as suitable for studying links between MASH and heart structural changes ahead of heart failure.

## Introduction

Metabolic dysfunction-associated steatotic liver disease (MASLD; formerly termed non-alcoholic fatty liver disease [NAFLD])[1] describes pathological liver conditions. lt is defined by the presence of hepatic steatosis (defined as fat accumulation in more than 5% of the hepatocytes),[2] accompanied by at least one cardiometabolic risk factor (elevated BMl, increased fasting glucose, hypertension, elevated plasma triglycerides and/or reduced HDL-cholesterol), and when further apparent causes (e.g. alcohol consumption) can be excluded.[1] Ongoing lobular inflammation (steatohepatitis) and hepatocyte injury (ballooning) drive disease progression to metabolic dysfunction-associated steatohepatitis (MASH; formerly termed non-alcoholic steatohepatitis [NASH])[1] and fibrosis. Eventually, it can develop into hepatocellular carcinoma (HCC), either directly or after excessive build-up of collagen deposits leading to cirrhosis and its cohort of life-threatening complications.[2,3] lt is estimated that MASLD affects approximately 30% of the general population.[4]

Case numbers of MASLD and cardiovascular diseases (CVDs) have been constantly rising in the last decades and represent a major burden for the public health systems.[5,6] lt has been demonstrated that patients suffering from MASLD (especially those with MASH and fibrosis) are at higher risk to develop CVD.[7] Actually more MASLD patients die from cardiovascular diseases (CVDs) than from liver-related events.[3,4] This association is grounded on metabolic risk factors such as abdominal obesity, hypertension, atherogenic dyslipidaemia and hyperglycaemia common to both CVD and MASLD. In addition, there is growing evidence that MASLD itself is an independent risk factor for CVD.[8] In human patients, MASLD not only promotes accelerated endothelial dysfunction and atherosclerosis [9] but also increases the risk of adverse cardiac remodelling (including hypertrophy)[10] that may progress to heart failure. However, the pathophysiological mechanisms underpinning this association are still poorly understood.[8,9] Processes currently debated in this context comprise disturbed lipid transportation, steatotic liver-triggered systemic exacerbation of oxidative stress and inflammation, dysregulated neuro-vascular control, genetic and epigenetic factors, as well as intestinal dysbiosis.[11] Although cardiovascular risk and MASH are linked epidemiologically and pathophysiologically, cardiovascular risk is, if any, rarely and poorly assessed in preclinical models and clinical trials. Hence, the major endpoint of clinical trials is liver fibrosis [12] while therapeutic effects on cardiovascular health are often overlooked.

The main aim of this study was to assess whether mice with MASH are prone to develop CVD and hence may serve as a model to explore the interplay and mechanism of a pathogenic axis linking MASLD and CVD. To this end, we used *foz/foz* mice (hereafter referred to as Foz) fed a high fat diet [11–15], and C57BL6/J mice fed a high fat diet with 30% fructose in the drinking water, as models for progressive fibrosing MASH. Foz mice carry a mutation in the *Almsl* gene. Patients with the Alstrom syndrome carry a mutation in that same gene and most patients develop a cardiomyopathy.[16] Foz mice are hyperphagic, they present early onset obesity and severe insulin resistance and complication thereof in multiple organs including the liver.[17] Yet, the cardiovascular system remains unexplored in these animals. We show that Foz mice with MASH develop cardiac hypertrophy with increased collagen deposition and activation of foetal genes in the heart. The cardiac function in these mice is still at a compensated level. However, tests with administration of angiotensin ll demonstrated increased susceptibility of these animals for decompensation. Our study demonstrates that both Foz and C57BL/6J mice are sound MASH models, with the former developing a phenotype more closely resembling the one observed in humans. The onset of a pathological cardiac phenotype occurred earlier and was more pronounced in Foz mice. We therefore consider the Foz mouse with MASH as a suitable model to address the liver/heart crosstalk in the early pathogenesis of CVD.

## Material & Methods

### Animals and diets

We used male non-obese diabetic (NOD.B10) fat aussie mice (Foz) bearing a homozygous truncating mutation in the *Almsl* gene and their wild type littermates (WT).[11,14]. In this study we restricted the investigation on male animals to avoid potential confounding effects of female hormones. As reported in both human patients and animal models a protective effect of oestrogen reduces the risk of developing MASLD [18] as well as CVD.[19] WT and Foz mice were fed either a standard rodent chow (termed normal diet [ND]; SAFE^®^ A03/13,5kcal% from fat/*SAFE SAS, France*) or a high fat diet (referred to as HFD; D12492/60 kcal% from fat/*Research Diets, USA*) starting at 5 weeks of age for 24 weeks (n=8-13/group) or 60 weeks (n=5-11/group). To assess cardiac adaptation to angiotensin ll overload, WT and Foz mice were fed a high fat diet for 10 weeks prior to subcutaneous implantation of osmotic mini pumps (Alzet, USA) to deliver vehicle (veh) or angiotensin ll (Angll) (at low dose: 0.2mg/kg/day; or at high dose: 1.4mg/kg/day) for 4 weeks during which the high fat diet was continued. WN and WH denote WT mice fed ND and HFD, respectively; likewise, FN and FH denote Foz mice fed ND and HFD, respectively. Furthermore, we used male C57BL/6J mice (Janvier labs, France) which were fed either a ND (serving as controls [CTRL]) or a HFD with additional fructose (30%) in the drinking water (denoted as HF+F), for a period of 60 weeks.

Animals had access to food and water ad libitum and were housed in a temperature and humidity-controlled environment under a day/night cycle with 12 hours each. The numbers of animals used for each experiment is mentioned in the description of the corresponding figures.

Animal experimentation followed regulatory guidelines for humane care of laboratory animals established at the Universite catholique de Louvain (UCLouvain) in compliance with European regulations. The study protocol was approved by UCLouvain’s ethics committee (2016/UCL/MD/003, 2020/UCL/MD/019 and 2022/UCL/MD/62). Experiments were performed in the Laboratory of Hepato-Gastroenterology (GAEN) and the Pole of Cardiovascular Research (CARD), respectively - both part of the Institute of Experimental and Clinical Research (lREC) at UCLouvain, Brussels, Belgium.

### Blood plasma analyses

Glucose (in whole blood; Accu Chek Aviva; Roche, Germany) and insulin (in blood plasma; Ultrasensitive Mouse Insulin ELlSA; Mercodia, Sweden) were measured after 4.5 hours of fasting (sampling from tail vein). Brain natriuretic peptide (Mouse BNP ELlSA; MyBioSource, USA) was measured in blood plasma obtained by cardiac puncture during sacrifice. Tests were performed according to manufacturer instructions.

### Sacrifice and tissue processing

Animals were terminally anesthetized with a ketamine (100mg/mL)/xylazine (20mg/mL) solution and the body weight (BW) was determined. The liver was excised, parts were fixed in 4% formalin at room temperature (RT) for 24h prior to paraffin embedding; additional specimens were snap frozen and kept at −80°C until further use. The heart was isolated and transferred into cold (4°C), modified (glucose-enriched, 11.5mM) Krebs-Henseleit-buffer, associated tissue was removed and the organ subsequently weighted. Tissue of the left ventricular (LV) wall was snap frozen and stored at −80°C for gene expression analyses. The lower half of the heart was fixed in 4% formalin (RT for 24h) and embedded in paraffin for histological and immunohistochemical analyses of transversal cross-sections.

### Liver and heart histology

Haematoxylin & eosin (H&E) staining of 5 µm thick sections of paraffin-embedded livers was used for routine histological evaluation and assessment of NAFLD activity score (NAS) according to Kleiner et al.,[20] as previously done for assessment of MASLD severity in this mouse strain;[13] a score higher than 5 diagnosed MASH. For immunohistochemical detection of F4/80, paraffin sections were treated with proteinase K, exposed to a primary rat anti-mouse F4/80 monoclonal antibody (1/200) (MCA497G, BioRad, USA). A rabbit anti-rat immunoglobulin (1/200) (Al-4001, Vector laboratories, USA) was then applied followed by a goat anti-rabbit streptavidin horseradish peroxidase-conjugated antibody (K4003 EnVision, Dako, USA). The peroxidase activity was revealed with diaminobenzidine (DAB, Dako, USA) and slides were counterstained with haematoxylin. In addition to score the extent of hepatic inflammation, these specimen were also used for identification of crown-like structures, a morphological characteristic in progression from simple steatosis to MASH.[21] Hepatic fibrosis was assessed on Sirius Red (SR)-stained liver sections, using QuPath software (versions 0.3.0 & 0.4.3, Dr. Peter Bankhead, Edinburgh, UK) to determine the total tissue collagen content.

Cardiomyocyte size was determined on wheat germ agglutinin (WGA)-stained, paraffin-embedded transversal sections of the heart. Picture analysis was performed with Visiopharm (VlS 2020.08 & 2023.01, Visiopharm, Denmark). The area of transversally cut cardiomyocytes was measured in multiple regions of the left ventricle. A minimum of 100 cells was analysed from each heart.

Cardiac fibrosis was analysed on SR-stained paraffin-embedded transversal sections of the heart. Myocardial fibrosis was quantified as SR-positive area relative to total parenchymal area using the QuPath software (versions 0.3.0 & 0.4.3, Dr. Peter Bankhead, Edinburgh, UK). Three sections separated by at least 100µm were analysed for each heart to prevent biased measurements due to fibrotic clusters.

### Gene expression

Total RNA was isolated from snap frozen LV wall or kidney after mechanical grinding (Precellys Evolution, Bertin Technologies, France) using the Maxwell RSC microRNA Tissue kit (Promega, USA). RNA was quantified via NanoDrop (Thermo Fisher Scientific, USA), reverse transcribed utilizing High Capacity cDNA Reverse Transcription Kit (Applied Biosystems, Lithuania). cDNA served as template for real-time quantitative PCR (RT-qPCR). Reactions were performed on a Rotor-Gene Q (Qiagen, Germany) using SYBR^TM^ Select Master Mix (Applied Biosystems, Lithuania) and primer pairs (Invitrogen, USA) listed in Table 1. Data were normalized to the expression of appropriate housekeeping genes (heart: *Gapdh;* kidney: *Ppia* (Table 1)) after invariance of this housekeeping gene was confirmed for all groups. n-fold expression was then calculated with the ΔΔCt-method using WN as reference for the other groups.

**Table 1:**
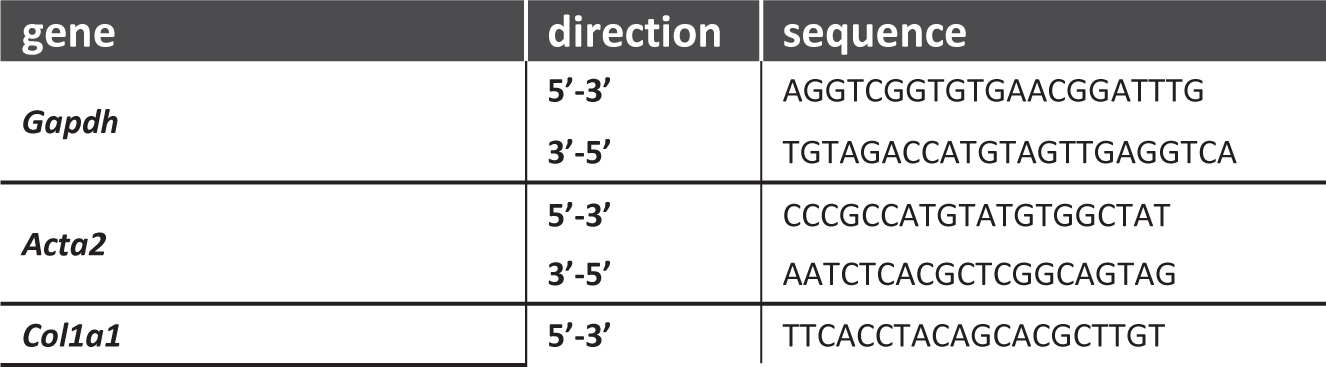
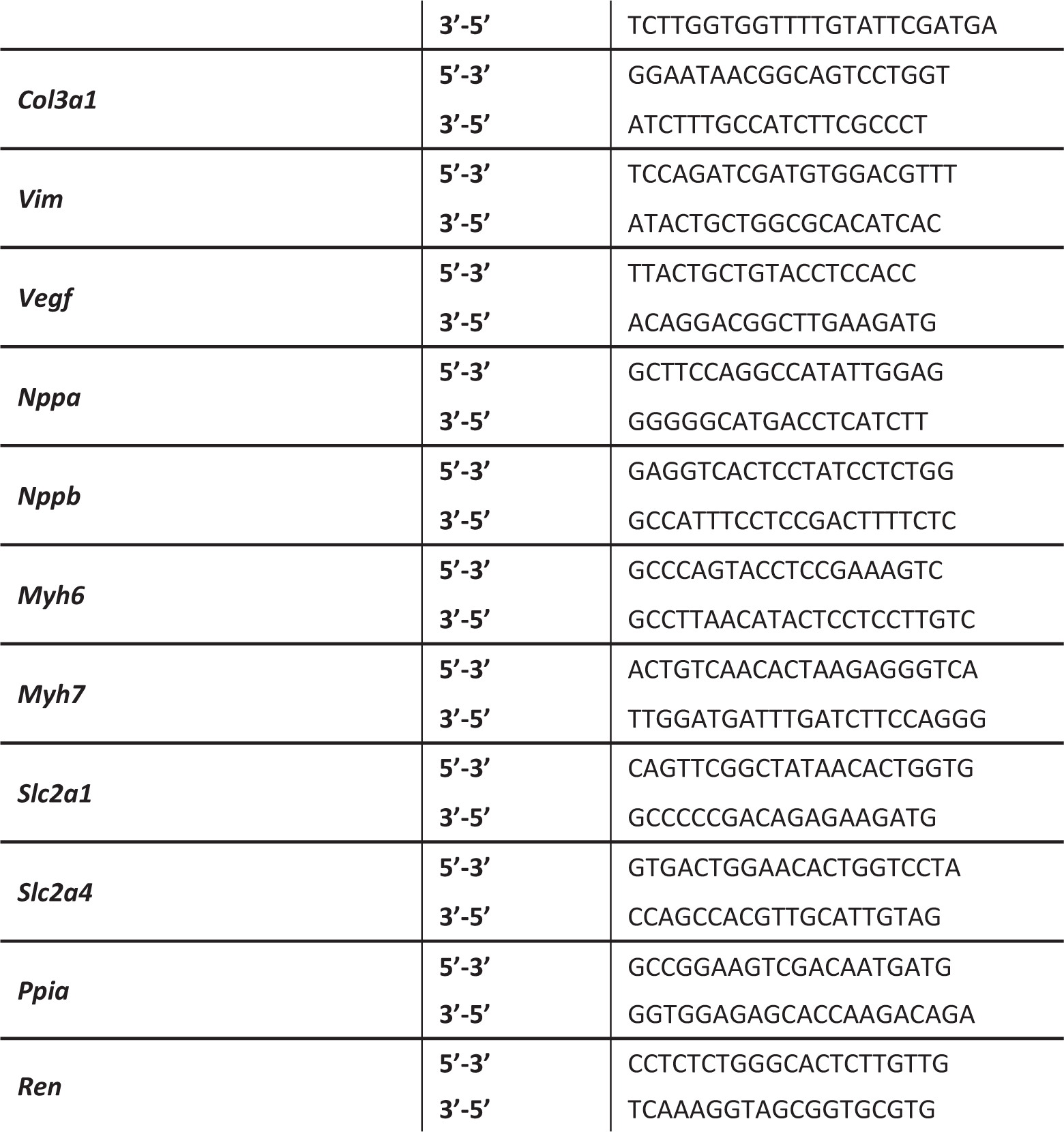
primers used for cardiac gene expression analysis via qPCR.

### Echocardiography

Transthoracic echocardiography was performed on a separate group of animals after 12 and 20 weeks of feeding. All procedures were performed on anesthetized mice (3-4% of isoflurane for induction and 1-2% for maintenance, in 100% oxygen) using a Vevo 2100 lmaging System (FUJlFlLM VisualSonics, Toronto, Canada) equipped with a 30 MHz transducer.

Systolic cardiac function parameters were measured as previously described [22] in 2-dimensional (2D) B-mode long axis and M-mode. Left ventricular volumes were measured using B-mode parasternal long-axis view, at end-systole and end-diastole, from which ejection fraction (EF %) was calculated. LV mass was calculated from LV long-axis measurements. Wall thickness dimensions (interventricular septum and posterior wall) was measured and fractional shortening (FS %) was calculated using internal LV dimensions measurements at end-systole and end-diastole, obtained from M-mode recordings.

Diastolic cardiac function parameters were calculated using doppler flow in an apical 4-chamber view, by applying pulsed wave doppler. E and A transmitral flow wave velocities were recorded at the tip of the mitral leaflet, parallel to the blood flow and the E/A ratio was calculated. *The* e’ mitral annular velocity was measured at the septal corner of the mitral annulus [22] and the E/e’ ratio was calculated.

All measurements and analyses were performed by the same experienced operator (EPD) who was blinded to the experimental groups.

### Pressure-volume loops

Pressure-volume (PV) loops were performed in Foz mice and their age-matched WT littermates after 24 weeks (WN, WH, FN, FH), or after 14 weeks (WH, FH; with and without Angll treatment). To complement the examination of cardiac function, cardiac ultrasound analyses (as described above) were performed on these animals in a separate session prior to the catheter measurements. The animals were kept anesthetized with isoflurane (1-1.5% in 100% of oxygen) and placed on a heating pad for the time of the procedure. Guided by high-resolution ultrasound-imaging a pressure-volume catheter (PVR-1045, Millar, USA), which was calibrated prior to each measurement, was introduced through the left carotid artery, advanced to the ascending aorta to measure systemic pressure and then inserted into the left ventricle by entering the left ventricle through the aortic valve. After an accommodation period of 15 minutes, the following basal cardiac parameters: HR (heart rate; bpm), EF (%), left ventricular end-diastolic pressure (LVEDP; mmHg), dP/dt max (mmHg/s), dP/dt min (mmHg/s), Tau (ms) were recorded for 5 minutes as well as during a series of at least 6 occlusions of the inferior vena cava (by exerting external pressure), to measure end-systolic and end-diastolic pressure volume relationship. Data recording was performed via LabChart pro software (version v8, ADInstruments, New Zealand). Determination of cardiac dimensions and guidance of catheter placement was performed with a high-resolution ultrasound system (Vevo 3100, 40 Mhz probe; FUJlFlLM VisualSonics, Toronto, Canada).

All measurements were performed by the same experienced operators (AM: PV loops; HE: echocardiography) who were blinded to the experimental groups and provided expertise for subsequent data analysis (performed by JL).

### Statistical analysis

Prior to evaluation each data set was tested for normality using the Shapiro-Wilk test.

Parametric data were analysed using two-way analysis of variance with subsequent Tukey’s or Dunnett’s multiple comparisons test or unpaired two-tailed *t*-test. Non-parametric data were evaluated using Kruskal-Wallis test with subsequent Dunn’s multiple comparisons test or unpaired two-tailed Mann-Whitney test. Data are depicted as mean ± standard error of mean. The utilized number of animals for each experiment is noted in the corresponding figure caption, with graphs displaying individual values.

All analyses were performed with GraphPad Prism 9.2.0 & 10.1.1 (GraphPad Software, Inc., USA). Differences were considered statistically different with a p-value < 0.05. The significance levels are displayed as follows: *p<0.05, **p<0.01, ***p<0.001, ****p<0.0001.

## Results

### FH mice, but not WT littermates, develop a severe MASH phenotype in a context of obesity and insulin resistance after 24 weeks feeding period

As previously reported,[23] Foz mice became rapidly obese when fed the HFD, though diet and genetic background concomitantly influenced body weight (Figure 1B). Foz mice fed the HFD also showed high fasting glycemia (Figure 1C) despite high fasting insulin concentration (Figure 1D) demonstrating severe insulin resistance. Insulin resistance was moderate in WT mice fed the HFD and in Foz mice on ND as supported by the levels of both insulin and glucose (Figure 1C/D).

**Figure 1:**
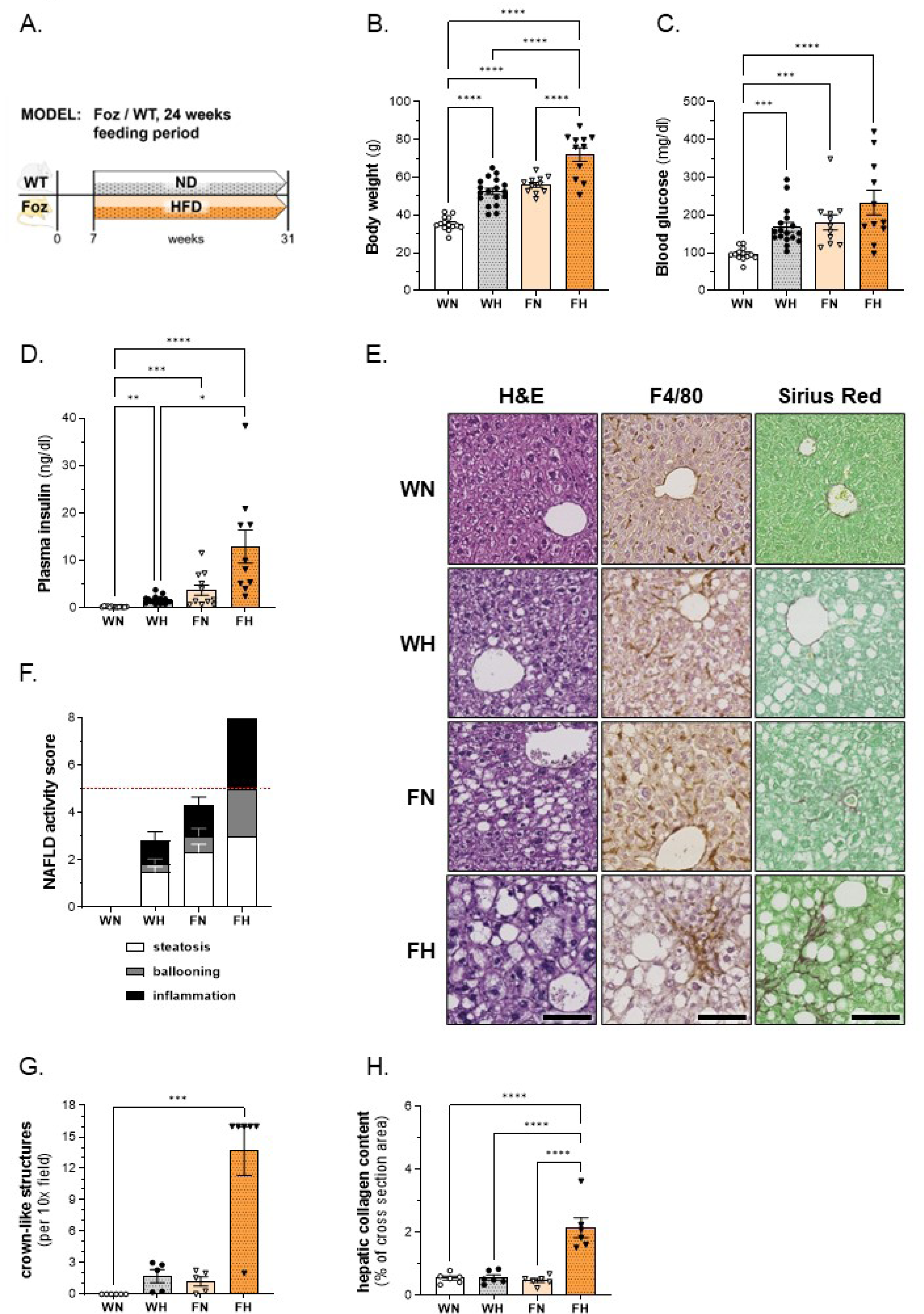
body weight, glucose homeostasis and liver phenotype in WT/Foz mice after 24 weeks feeding period. (A) experimental setup; (B) BW; (C) glycemia and (D) insulinemia after 4.5h of fasting; (E) representative histological pictures of H&E-, F4/80- and SR-stained sections, scale bar: 100µm; (F) NAS based on evaluation of steatosis (white bars), ballooning (grey bars) and inflammation (black bars), threshold for MASH displayed as dashed red line; (G) number of crown-like structures per 10x field of view (using aforementioned F4/80-stained sections); (H) quantification of hepatic collagen content in % of total area. **Animals: n=G-l8 per group; Statistics: (B, H): two-way ANOVA with Tukey’s multiple comparisons test; (C, D, G): Kruskal-Wallis with Dunn’s multiple comparisons test.**

As expected,[24] FH mice had an enlarged liver that exhibited severe panlobular steatosis with prominent inflammation and ballooning (Figure 1E). By contrast, the liver was normal in WN, while WH and FN livers presented an intermediate phenotype with moderate steatosis, mild inflammation, and inconstant ballooning. Hence, when histological sections were scored for steatosis, ballooning and inflammation, the assigned NAFLD activity score (NAS) was maximal in FH (for all three parameters in each animal), intermediate in FN and WH and null in WN (Figure 1F). lHC with the macrophage marker F4/80 supported a moderate activation of liver macrophages in the WH and FN groups compared to controls, but a massive infiltration with formation of aggregates and crown-like structures was observed in FH (Figure 1E). In line, inflammatory foci were absent in WN, rare in WH and FN and markedly increased in numbers in FH (Figure 1G). As shown on SR-stained sections, fibrosis was absent in WN, minimal collagen deposition was seen in WH and was slightly more pronounced in FN, while MASH in FH was associated with diffuse lobular pericellular fibrosis typically seen in human MASH (Figure 1E). The quantification of the hepatic collagen content confirmed significant fibrosis in FH (Figure 1H).

Thus, mice in our 24 weeks-cohort encompass the spectrum of MASLD and metabolic syndrome: FH mice are overtly obese, insulin resistant and exhibit severe fibrosing MASH whereas WH and FN are moderately obese and insulin resistant and exhibit steatosis with moderate levels of inflammation and cell injury but inconspicuous fibrosis. WN have a normal liver.

### FH mice show LV hypertrophy with adverse remodelling after 24 weeks of HFD

After 24 weeks of HFD diet, heart weight/tibia length ratios were significantly higher in Foz mice than in WT mice independently of diet (Figure 2B). The mean cardiomyocyte size was larger in FH compared to WN mice, whereas the average size of WH and FN cardiomyocytes was between those of the aforementioned groups (Figure 2C, D). Increased collagen deposition on SR-stained sections was confirmed by morphometrical quantification showing higher myocardial collagen content in Foz mice with MASH compared to WN control mice (Figure 2E, F).

**Figure 2:**
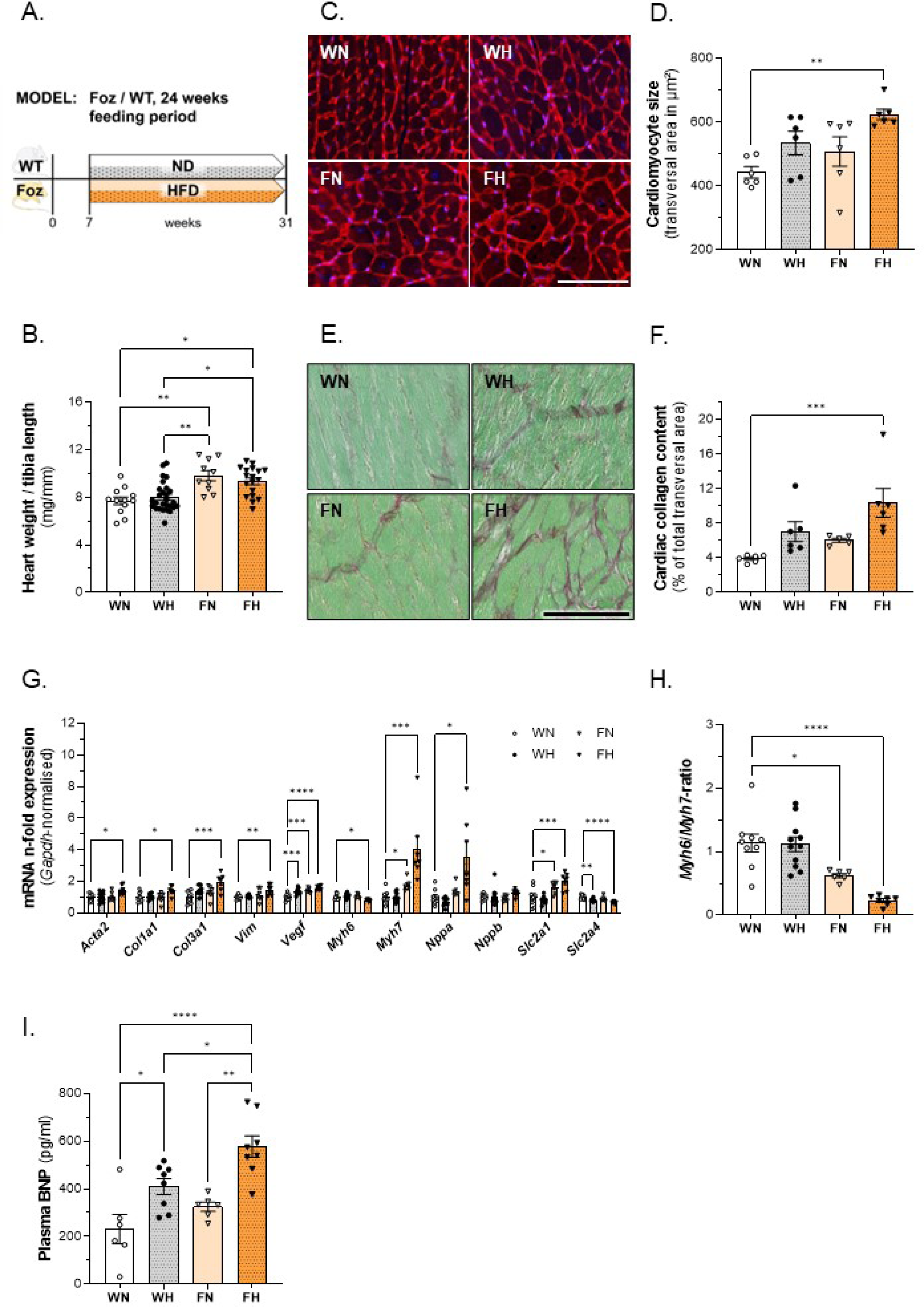
characterization of cardiac phenotype in WT/Foz mice after 24 weeks feeding period. (A) experimental setup; (B) heart weight normalized with tibia length; (C) representative WGA-staining of cardiomyocytes for determination of cell size, scale bar = 100µm; (D) average transversal cardiomyocyte size; (E) representative Sirius red staining to quantify cardiac collagen content; (F) average collagen content from cardiac transversal cross-sections of three different cuts (distance;?100µm); (G) cardiac gene expression levels normalized with *Gapdh*, n-fold expression calculated with ΔΔCt-method using WN as reference for the other groups; (H) *Myh6*/*Myh7* ratio using WN as reference for the other groups; (l) plasma BNP levels. **Animals: n=G-27 per group; Statistics: (D, G, H, I): one-way ANOVA with Tukey’s or Dunnett’s (only for G) multiple comparisons test; (B, F, G): Kruskal-Wallis with Dunn’s multiple comparisons test.**

We evaluated the level of expression of genes associated with adverse LV remodelling. Compared to WN mice, *collagen I (Collal)* and *III (Col3al)*, *alpha smooth muscle actin (Acta2)* and *vimentin (Vim)* mRNA were upregulated in FH mice after 24 weeks of diet (Figure 2G). *Vascular endothelial growth factor (Vegf)* expression was significantly higher in FN compared to WN mice and even more in FH. Expression of the foetal myosin heavy chain 7 (*Myh7)* was upregulated while that of adult *Myh6* was downregulated in FH hearts. Hence the *Myh6/Myh7*-ratio was the lowest in the FH group (Figure 2H). This re-induction of the foetal gene *Myh7*, encoding for β-myosin heavy chain (β-MHC), is considered a hallmark of pathological hypertrophy.[25] Regarding glucose transporters, we observed the upregulation of solute carrier family 2 member a 1 (*Slc2al; Glutl*) accompanied by concomitant downregulation of *Slc2a4* (*Glut4*) in FH hearts supporting insulin resistance and increased unregulated glucose uptake in the face of permanent hyperglycaemia in the remodelled myocardium of FH. mRNA of natriuretic peptide A (*Nppa*) but not B (*Nppb*) was upregulated in FH hearts, while BNP serum levels were significantly higher in FH than in WN controls, here again confirming pathological cardiac hypertrophy (Figure 2l).

Taken together, our data show evidence of pathological cardiac remodelling in Foz mice with MASH after 24 weeks feeding period. This phenotype was not seen in WT animals fed the HFD over the same period.

### Cardiac function is not compromised in Foz mice with MASH

In the FH group, a significant thickening of the left ventricular wall and the septum was systematically by echocardiography observed after 20 weeks of feeding (Supplements, table 1). These differences were already significant after 12 weeks feeding period. The computed LV mass also tended to be higher in FH compared to other groups. The LV diameter at end diastole was also larger in FH after 20 weeks of feeding but not after 12 weeks. lmportantly, LV systolic function parameters such as ejection fraction, fractional shortening, and stroke volume, were comparable to those of WN controls at both time points. We used Doppler echocardiography to further evaluate diastolic function and filling pressure of the left ventricle (E/A-ratio and E/e’-ratio).[26] After 12 weeks, there were no significant differences among the groups; unfortunately, we could not acquire doppler data in the FH group after 20 weeks, as the mice were far too obese (63.78 ± 1.49 g) (Supplements, table 1). Our attempts to obtain proper data regarding left atrial area were not successful in any group at any time point. To counterbalance this data gap, we measured cardiac haemodynamics in a separate cohort of mice using a Millar catheter, the “gold standard” method to study cardiac pathophysiology and haemodynamics, particularly diastolic function (representative PV loops are displayed in Supplements figure 1). Before proceeding with catheterization, we confirmed that ejection fraction was similar in all groups (Figure 3B) and comparable to the initial cohort (Supplements, table 1). Haemodynamic measurements obtained by PV loops showed that LV end-systolic pressure was among groups (Figure 3C). The end-diastolic pressure showed a tendency to increase in WH mice, but no significant changes were found between the groups (Figure 3D). There was also no difference in the changes in the pressure/volume relationship during vena cava occlusion between the four groups, whether at the end of the systole (Figure 3E) or at the end of the diastole (Figure 3F). The measured values for maximum (Figure 3G) and minimum (Figure 3H) rate of pressure change in the left ventricle exhibited no significant differences between the groups. And both the end-systolic (Figure 3l) and -diastolic volume (Figure 3J) showed similar values for WN and FH. Hence the data support that in Foz mice with MASH there was cardiac hypertrophy without cardiac dysfunction.

**Figure 3:**
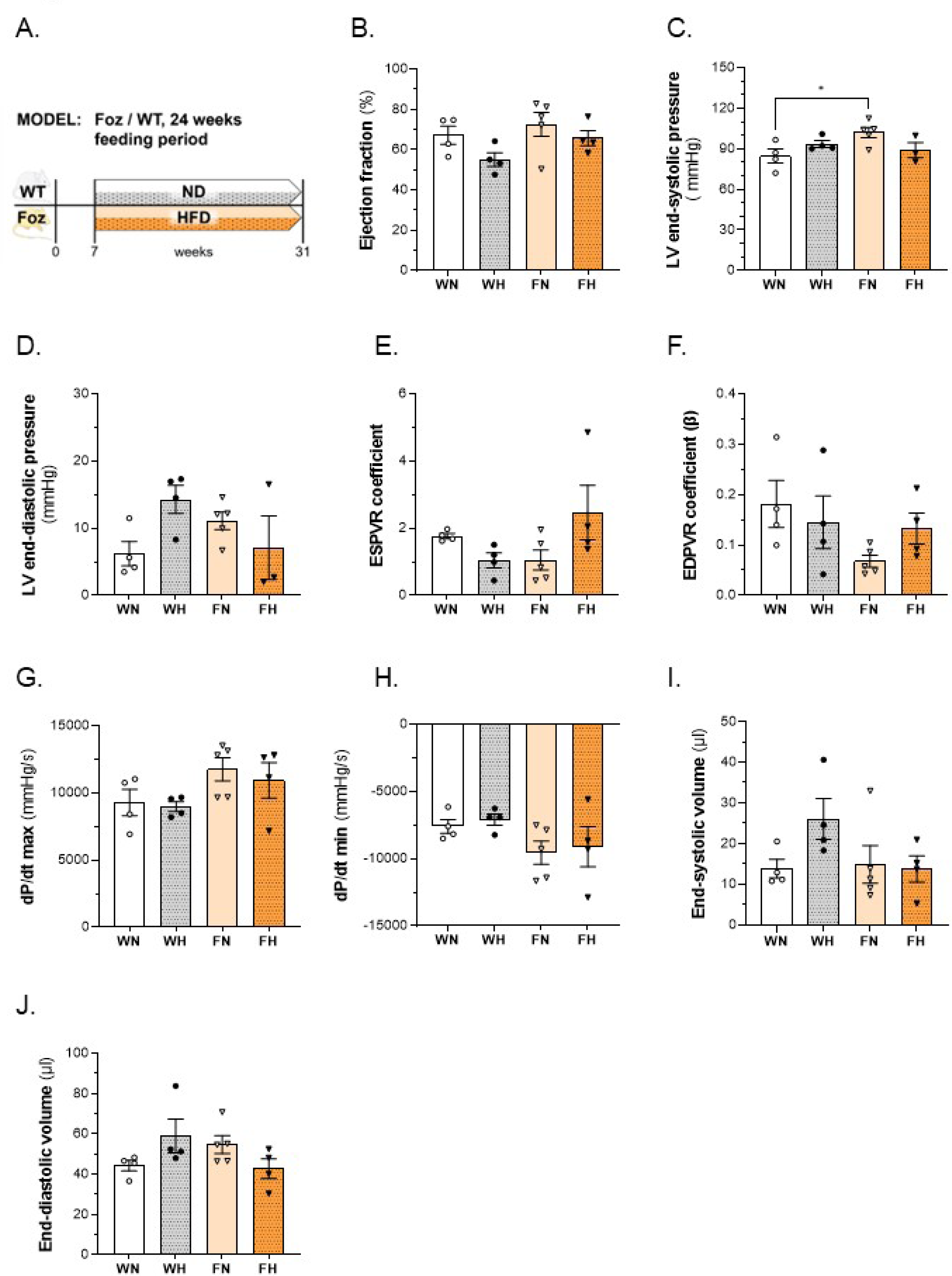
assessment of cardiac function in WT/Foz mice after 24 weeks feeding period. (A) experimental setup; (B) cardiac ejection fraction; (C) pressure in the left ventricle at the end of systole; (D) pressure in the left ventricle at the end of diastole; (E) pressure/volume relationship during vena cava occlusion at the end of systole; (F) pressure/volume relationship during vena cava occlusion at the end of diastole; (G) maximum rate of pressure change in the left ventricle; (H) minimum rate of pressure change in the left ventricle; (l) left ventricular volume at the end of systole; (J) left ventricular volume at the end of diastole. **Animals: n=2-8 per group; Statistics: (B, C, D, E, F, G; H, I, J): two-way ANOVA with Tukey’s multiple comparisons test.**

### Foz are more sensitive to exogenous Angii

We challenged FH with Angll to test whether increase in the post-charge would precipitate adverse remodelling or heart failure in mice with MASH. We administered angiotensin ll or vehicle via osmotic minipump during the last 4 weeks of a 14 weeks HFD regimen. At this time point FH mice are known to have established MASH.[13]

We first treated the mice with a small dose of angiotensin ll (0.2 mg/kg/day) chosen to not increase blood pressure but to trigger diastolic dysfunction.[27] Indeed, we observed neither change in blood pressure (BP), nor activation of a negative feed-back on renin expression. At this dosage, no effect was observed on the heart (not shown). We then used Angll at the dose of 1.4 mg/kg/day. Measurements of systolic (Figure 4B) and diastolic (Figure 4C) blood pressure exhibited similar and significant elevation in WT and Foz animals treated with Angll; values from the untreated control groups were similar to each other, too. Angll reduced kidney renin mRNA expression levels in WT mice, however in Foz mice this decrease was less pronounced and not significant (Figure 4D). The body weight was not altered by Ang ll in either group compared to the corresponding control group (Figure 4E). The same applies to the relative heart weight, on which Angll administration had no effect compared to vehicle-treatment (Figure 4F). We complemented these analyses by measurements of the cardiomyocyte size. In accordance with the observed heart weight, the cardiac muscle cells were not affected by Angll-administration (Figure 4G). We also investigated the direct impact of the Angll-treatment and the resulting increased cardiac workload on genes associated with adverse remodelling in the left ventricular wall. Figure 4H reports the fold change in mRNA expression in animals treated with Angll in comparison to those treated with vehicle. In general, Foz mice demonstrated to be more affected by Angll-administration than WT mice. *Collal* and *Col3al* showed moderately but significantly elevated expression levels in these animals, whereas major upregulation was detected for *Myh7* and *Nppa* (Figure 4H). As a result, the *Myh6/7* ratio was more decreased upon Angll treatment compared to vehicle controls in Foz than in WT mice (Figure 4l).

**Figure 4:**
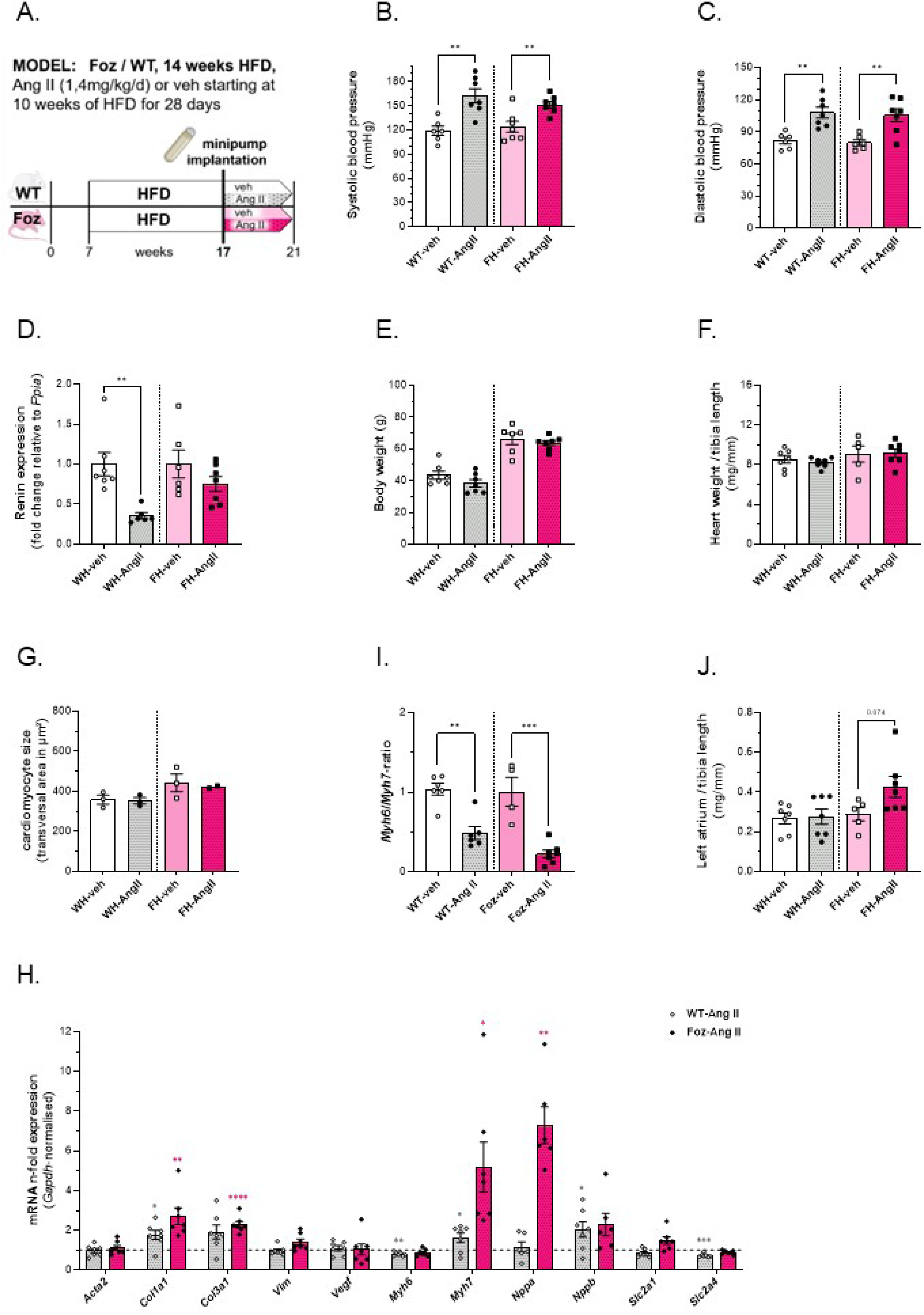
impact of 4 weeks of AngII-treatment on nephrotic renin expression, body weight and cardiovascular phenotype in WT/Foz mice on HFD. (A) experimental setup; (B) blood pressure at the end of systole; (C) blood pressure at the end of diastole; (D) renin expression in the kidney, normalized with *Ppia*, n-fold expression calculated with ΔΔCt-method using veh as reference for the corresponding Angll-group; (E) body weight; (F) heart weight normalized with tibia length; (G) average transversal cardiomyocyte size; (H) cardiac gene expression levels normalized with *Gapdh*, n-fold expression calculated with ΔΔCt-method using veh as reference for the corresponding Angll-group (asterics for WT: grey; asterics for Foz: pink); (l) *Myh6*/*Myh7* ratio, (J) left atrial weight normalized with tibia length. **Animals: n=2-7 per group; Statistics: (B, C, D, E, F, G, H, I, J): unpaired two-tailed t-test; (D, H, I): unpaired two-tailed Mann-Whitney test.**

Although Foz mice showed a weaker effect of renin-expression in the kidney compatible with some resistance to the action of angiotensin ll at that level (see Figure 4D), the treatment with Angll concurrently demonstrated a more pronounced impact on cardiac genes involved in adverse remodelling of the heart.

To complement our findings regarding the impact of Angll on the cardiac phenotype in Foz mice, the cardiac function was examined, too. No significant differences were observed for ejection fraction (Figure 5B) and fractional shortening (Figure 5C) of the different groups. The left ventricular mass, however, was significantly increased in Angll-treated FH but not WH mice, compared to vehicle controls (Figure 5D). Using a catheter, we then measured left ventricular pressure and volume. FH mice treated with Angll showed significantly elevated LV end-systolic pressure (Figure 5E) and a tendency towards increased LV end-diastolic pressure (Figure 5F). In comparison to their vehicle control group. The measured maximal rise of left ventricular pressure (dP/dt max), a parameter used for assessment of myocardial contractility, showed no significant differences between the treatment- and control groups (Figure 5G). This was also the case for the maximal pressure drop rate (dP/dt min), which is utilized to evaluate the ventricular capability to relax during diastole. Interestingly, although not significant, Foz mice tended to have a slower pressure drop rate when treated with Angll compared to vehicle, whereas the WT groups exhibited no such pattern (Figure 5H). In comparison to their vehicle-treated controls, the end-systolic volume showed a strong tendency (p=0.052) to be declined in the FH group to which Angll was administered (Figure 5l), whereas the end-diastolic volume was significantly reduced (Figure 5J) in those mice. Except for WH treated with vehicle, mice in the other groups did not tolerate for long the retrograde catheterization through the aortic valve (acute aortic insufficiency), we were therefore not able to perform the vena cava occlusion in a sufficient number of animals. To complement these analyses, we additionally determined the relative left atrial weight as an indicator for atrial remodelling. Our data show that tended to be higher (p=0.074) in Angll-treated FH mice (Figure 4J) compared to Foz-veh mice.

**Figure 5:**
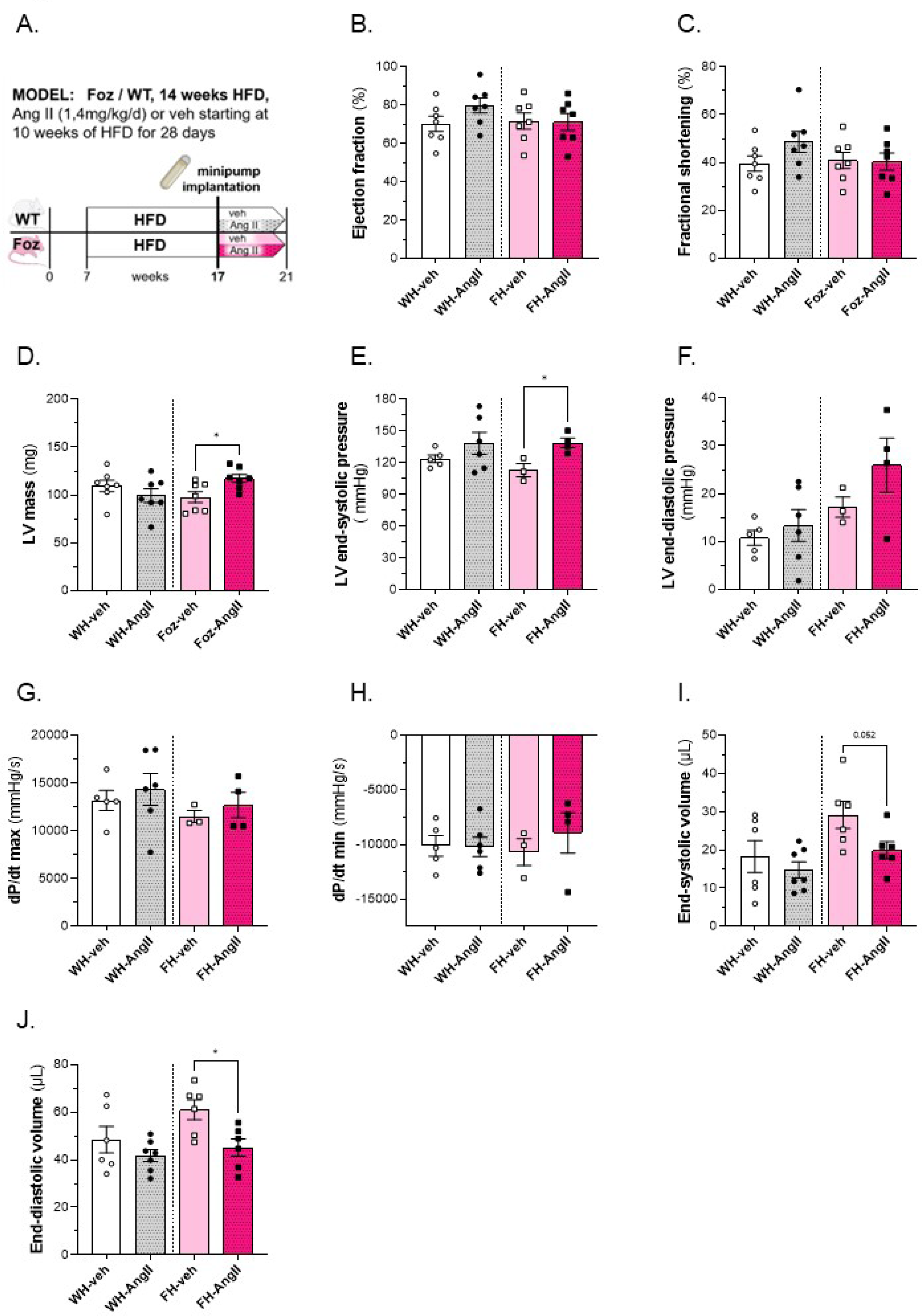
impact of 4 weeks of AngII treatment on cardiac function in WT/Foz mice on HFD. (A) experimental setup; (B) cardiac ejection fraction; (C) fractional shortening; (D) left ventricular mass; (E) basal pressure in the left ventricle at the end of systole; (F) basal pressure in the left ventricle at the end of diastole; (G) maximum rate of pressure change in the left ventricle; (H) minimum rate of pressure change in the left ventricle; (l) left ventricular volume at the end of systole; (J) left ventricular volume at the end of diastole. **Animals: n=3-7 per group. Statistics: (B, C, D, E, F, G, H): unpaired two-tailed t-test; (C, D, E): unpaired two-tailed Mann-Whitney test.**

Taken together, the cardiac function was largely compatible in all four groups, though Angll-administration decreased the end-diastolic volume and increased LV mass in Foz mice compared to vehicle-treatment. Thus, angiotensin amplifies the transcriptomic signature of adverse cardiac remodelling in FH mice with non-significant impact on cardiac function but with a deleterious development towards diastolic dysfunction.

### Aged Foz mice exhibit aggravation of hepatic but not cardiac pathology

To evaluate whether the duration of metabolic disorder and severity of MASH would aggravate the cardiac phenotype, we analysed WT and Foz mice after 60 weeks on HFD or regular rodent chow. Despite being obese (Figure 6B), the insulin sensitivity has improved in old FH mice as they exhibited a normal glycemia (Figure 6C), unlike their younger counterparts (Figure 1C) tough in the face of elevated serum insulin levels (Figure 6D). At the histological level, except for WN controls, all mice presented some levels of pericentrilobular damage, chronic inflammation, and variable fibrosis in the absence of significant steatosis or ballooning (Figure 6E). Lack of these two essential criteria prevented proper scoring to create a NAS for this experimental approach. The severity of the damage increased from mild in WH, to moderate in FN and severe in FH. As in the 24 weeks cohort, the number of crown-like structures after 60 weeks was still highest in HFD-fed Foz mice compared to the other groups (Figure 6F), however the total count was much lower. In FH pericentral bridging fibrosis was observed. The histological aspect is compatible with a prolonged exposure to (oxidative) stress and centrilobular necrosis with post necrotic remodelling and fibrosis or burn out MASH (Figure 6G). Histological assessment also revealed the development of liver cancer in half of the animals fed a HFD (WH: 1 in 2 mice; FH: 2 in 4 mice). Compared to 24wks-fed counter parts, we observed no further aggravation of the cardiac hypertrophy (Figure 7B), cardiomyocyte hypertrophy (Figure 7C, D) or fibrosis (Figure 7E, F) in FH after 60 weeks. In fact, the average cardiac collagen content as well as the cardiomyocyte size of aged FH mice had decreased in comparison to the younger FH group (Figures 2E, F and C, D, respectively). At this more advanced age, WN had developed cardiac fibrosis and there was no difference in the amount of fibrillar collagen between groups (Figure 7E, F). Similarly, at the gene expression level, changes were of lesser magnitude in FH after 60 weeks (Figure 7G) than after 24 weeks compared to age matched controls (Figure 2G). Furthermore, *Myh6/Myh7*-ratios were comparable in all four groups, no significant differences were found (Figure 7H). Plasma BNP levels were similar in WH and FH mice and significantly elevated compared to WN animals (Figure 7l). There was no difference in lung weight between the groups and therefore no lung oedema in HFD-fed Foz mice (Figure 7J).

**Figure 6:**
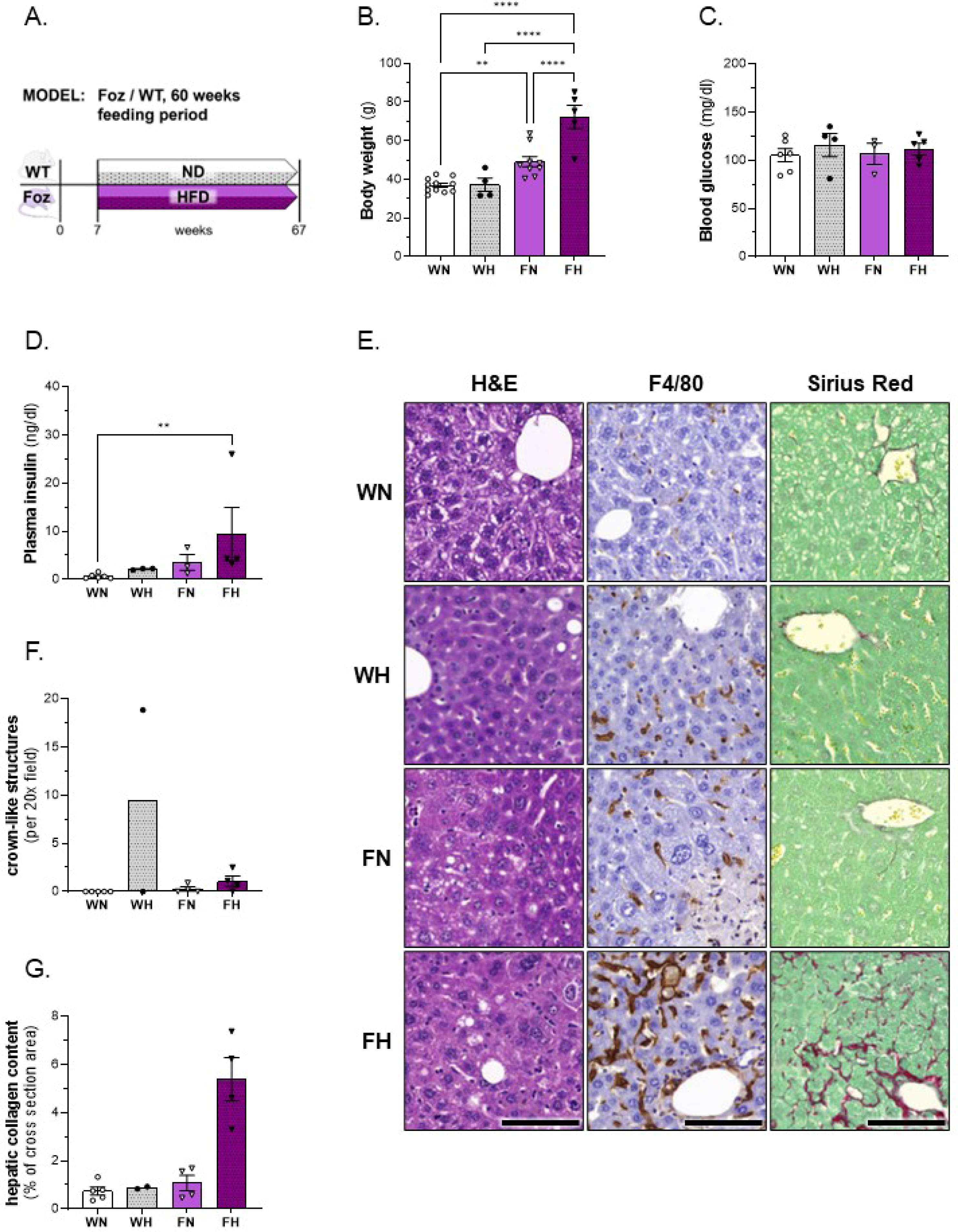
body weight, glucose homeostasis and liver phenotype in WT/Foz mice after G0 weeks feeding period. (A) experimental setup; (B) BW; (C) glycemia and (D) insulinemia after 4.5h of fasting; (E) representative histological pictures of H&E-, F4/80- and SR-stained sections, scale bar: 100µm; (E) Quantification of CPA in %; (F) number of crown-like structures per 20x field of view (using aforementioned F4/80-stained sections); (G) quantification of hepatic collagen content in % of total area. **Animals: n=2-ll per group; Statistics: (B, C): two-way ANOVA with Tukey’s multiple comparisons test; (D): Kruskal-Wallis with Dunn’s multiple comparisons test.**

**Figure 7:**
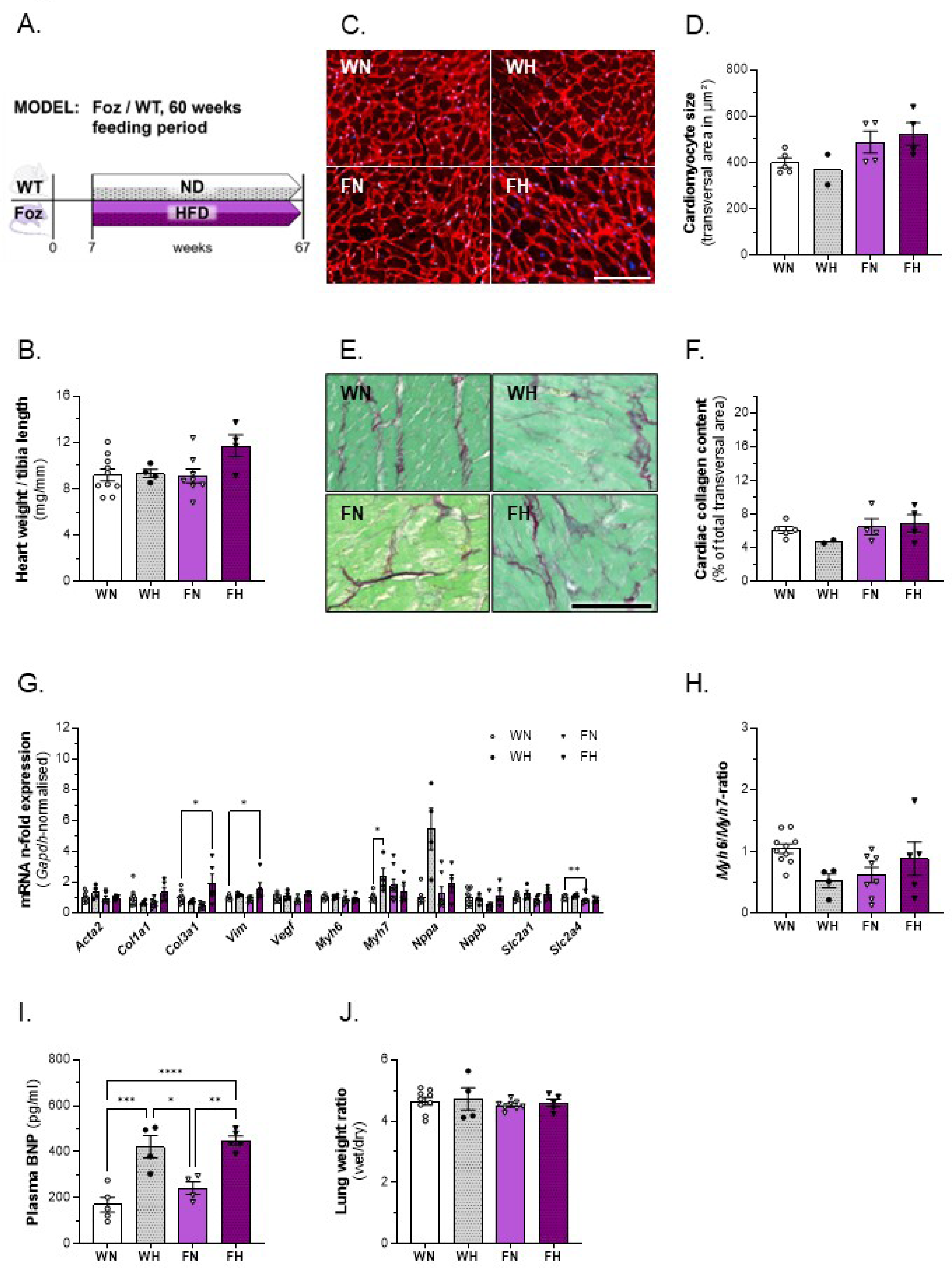
characterization of cardiac phenotype, plasma BNP and lung weight in WT/Foz mice after G0 weeks feeding period. (A) experimental setup; (B) heart weight normalized with tibia length; (C) representative WGA-staining of cardiomyocytes for determination of cell size, scale bar = 100µm; (D) average transversal cardiomyocyte size; (E) representative SR-staining to quantify cardiac collagen content; (F) average total collagen content from cardiac transversal cross-sections of three different cuts (distance;?100µm); (G) cardiac gene expression levels normalized with *Gapdh*, n-fold expression calculated with ΔΔCt-method using WN as reference for the other groups; (H) *Myh6*/*Myh7* ratio using WN as reference for the other groups; (l) plasma BNP levels; (J) lung wet weight to dry weight ratio. **Animals: n=2-l0 per group; Statistics: (B, G, H, I, J): one-way ANOVA with Tukey’s or Dunnett’s (only for G) multiple comparisons test.**

Thus, prolonged nutritional overload, metabolic and hepatic disorders do not aggravate the cardiac phenotype nor cause the decompensation of the cardiomyopathy. This was at variance with hepatic centrilobular damage and sinusoid dilations, compatible with an increased filling pressure of the right ventricle.

### Aged C57BL/6J mice with MASH show adverse cardiac remodelling without developing cardiac hypertrophy

To verify the results we found in Foz mice in an independent model, we analysed C57BL6/J mice that received a high fat diet and fructose-rich beverage for 60 weeks. At this stage the mice on HF+F are obese (Figure 8B) and their blood glucose (Figure 8C) and insulin (Figure 8D) levels are elevated compared to those of the ND-fed control group, characteristics of insulin resistance. Furthermore, they have developed severe MASH (Figure 8E, F) with significant inflammation (Figure 8G) and fibrosis (Figure 8H). Though they have significantly increased relative cardiac weight (Figure 9B), we did not observe cardiomyocyte hypertrophy (Figure 9C, D). However, cardiac fibrosis levels were significantly increased in HF+F mice (Figure 9E, F) compared to age-matched controls. These findings are in accordance with alterations at the gene expression level, where upregulation of *Collal* and *Col3al*, *Vimentin*, *Vegf* and *Slc2al* (Figure 9G) as well as downregulation of *Myh6* suggest the operation of cardiac remodelling. Although not significantly lowered (p=0,057), a reduced *Myh6/Myh7* ratio in FH+H mice (Figure 9H) is in accordance with the results of the gene expression analysis. Interestingly, plasma BNP levels were significantly increased in FH+F mice compared to the control group (Figure 9l). Determination of the lung wet/dry weight ratio showed slightly but significantly decreased levels in HF+F mice compared to the control group (Figure 9J); an observation that contradicts the presence of pulmonary oedema. However, in the animals suffering from MASH we observed a significant enlargement of the left atrium (Figure 9K), an indicator for ongoing atrial remodelling.

**Figure 8:**
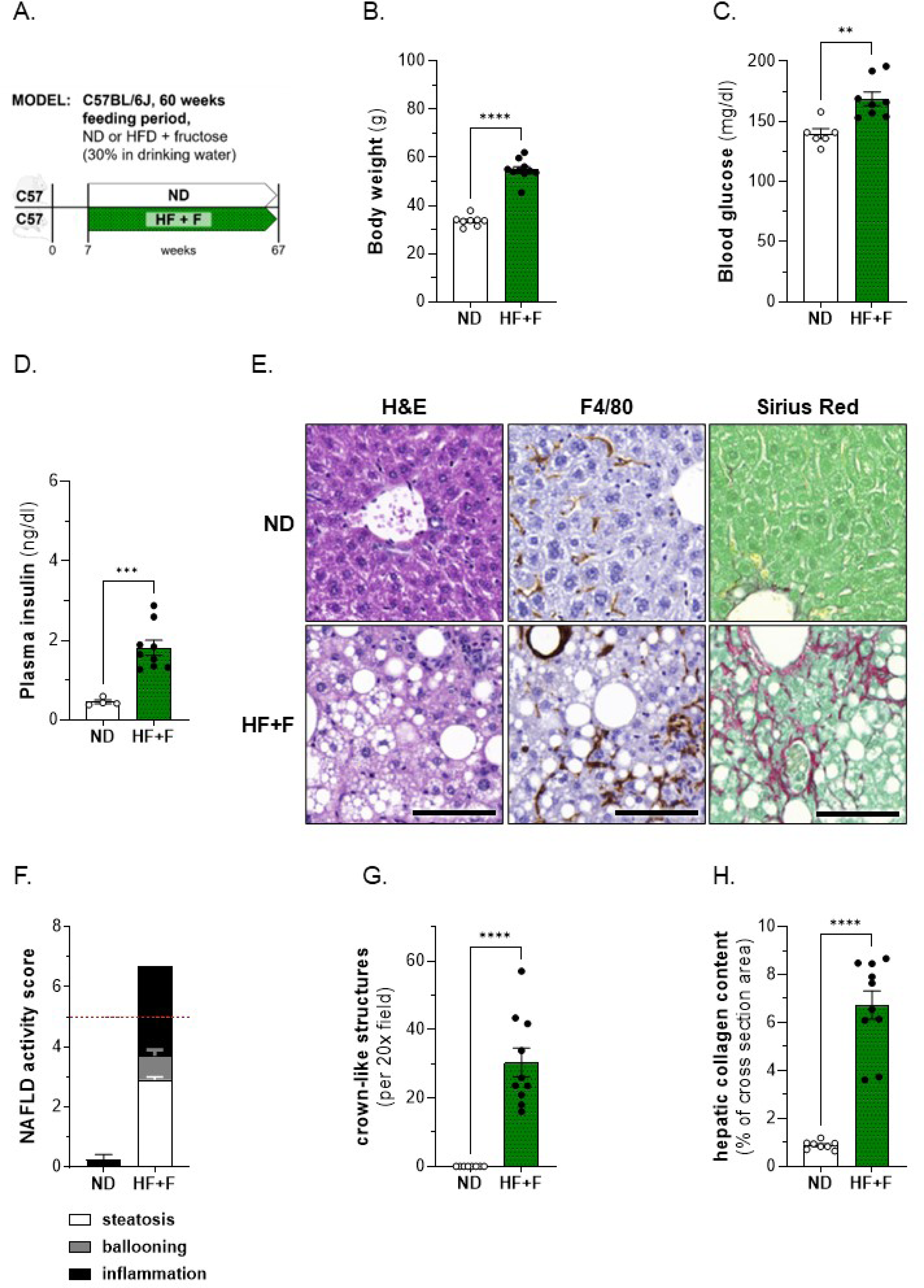
body weight, glucose homeostasis and liver phenotype in CS7BL/GJ mice after G0 weeks feeding period. (A) experimental setup; (B) BW; (C) glycemia and (D) insulinemia after 4.5h of fasting; (E) representative histological pictures of H&E-, F4/80- and SR-stained sections, scale bar: 100µm; (F) NAS based on evaluation of steatosis (white bars), ballooning (grey bars) and inflammation (black bars), threshold for MASH displayed as dashed red line; (G) number of crown-like structures per 20x field of view (using aforementioned F4/80-stained sections); (H) quantification of hepatic collagen content in % of total area. **Animals: n=4-l0 per group; Statistics: (B, C, D, H): unpaired two-tailed t-test.**

**Figure 9:**
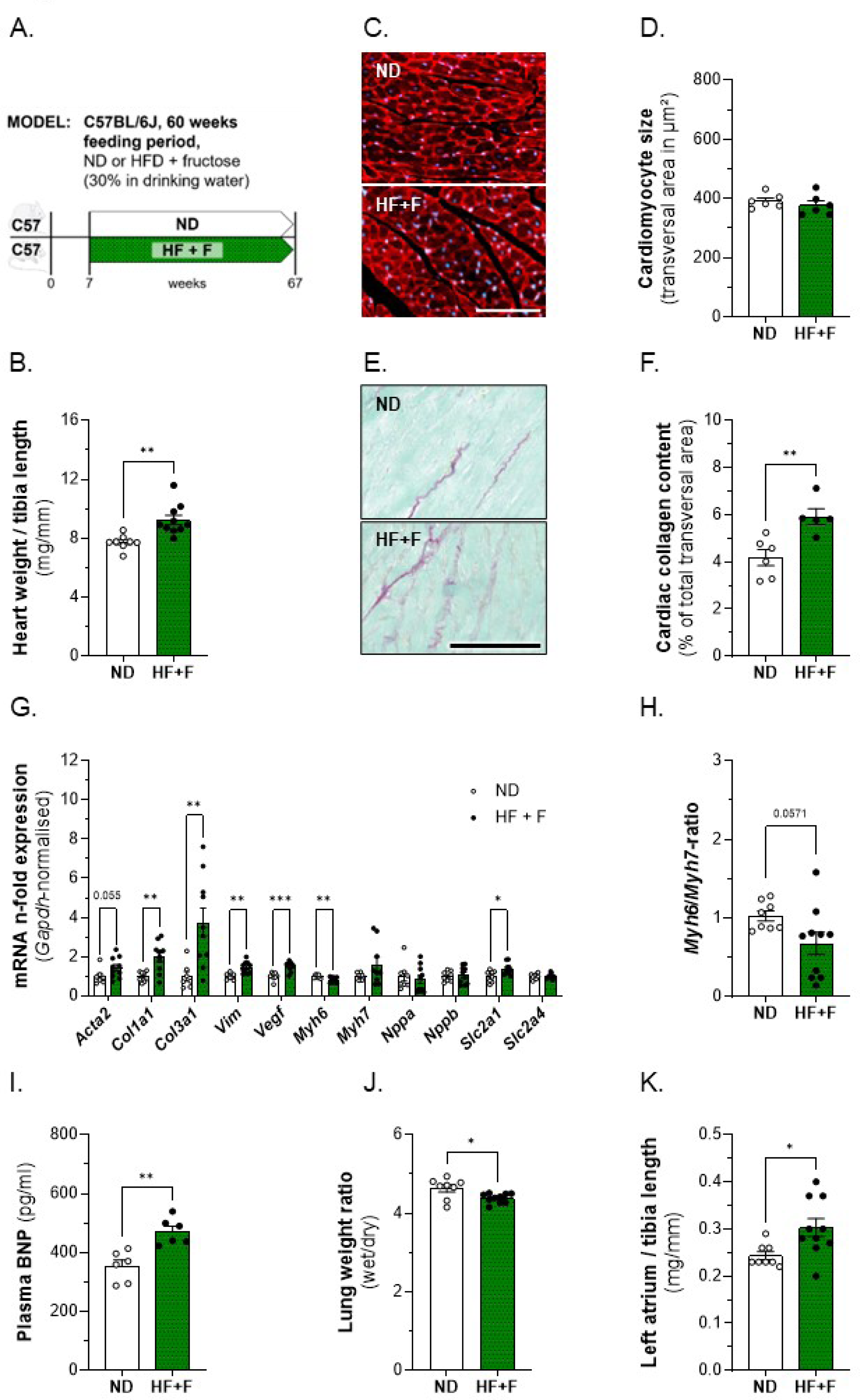
characterization of cardiac phenotype, plasma BNP and lung weight in CS7BL/GJ mice after G0 weeks feeding period. (A) experimental setup; (B) heart weight normalized with tibia length; (C) representative WGA-staining of cardiomyocytes for determination of cell size, scale bar = 100µm; (D) average transversal cardiomyocyte size; (E) representative SR-staining to quantify cardiac collagen content; (F) average total collagen content from cardiac transversal cross-sections of three different cuts (distance;?100µm); (G) cardiac gene expression levels normalized with *Gapdh*, n-fold expression calculated with ΔΔCt-method using ND as reference for the HF+F group; (H) *Myh6*/*Myh7* ratio using ND as reference for HF+F; (l) plasma BNP levels; (J) lung wet weight to dry weight ratio, (K) left atrial weight normalized with tibia length. **Animals: n=S-l0 per group. Statistics: (B, D, F, G, H, I, J): unpaired two-tailed t-test; (G, K) unpaired two-tailed Mann-Whitney test.**

All in all, the mice fed HF+F for 60 weeks had developed severe MASH with bridging fibrosis; although no cardiac hypertrophy was detected, our findings suggest that the observed changes are not simple physiological remodelling but signs of adverse alterations in the hearts of these animals.

## Discussion

In this work we describe for the first time that HFD-fed Foz mice with MASH, a faithful model of progressive human MASLD, exhibit cardiac hypertrophy, fibrosis, and dysregulation of genes involved in pathological structural changes in the heart. However, such an adverse cardiac remodelling, even when further aggravated by long term exogenous angiotensin ll administration, does not result in major alteration of cardiac function. Cardiac remodelling was also associated with severe MASH in C57BL6/J mice fed a fat- and fructose-rich regimen. Our data support that MASH, and not simply the high calory diet, causes accelerated aging of the heart.

CVD and MASLD are affecting a growing number of people worldwide. Data from human patients indicate that MASLD is an additional and independent risk factor for cardiovascular events.[28,29] Despite sharing several risk factors (namely obesity, T2DM, dyslipidaemia),[30] CVD and MASLD are still considered separately in clinical trials, with the therapeutic end-point for liver targeted therapy being MASH resolution and decreased liver fibrosis.[12] A potential beneficial impact on the cardiovascular system as a positive “side-effect” of eliminating MASH in the liver is thereby disregarded. We show that mice developing fibrosing MASH, but not those with benign steatosis or receiving a fat-rich diet, develop a cardiac phenotype with adverse structural but not functional changes. The pathological alterations affect cardiac gene expression, accumulation of extracellular fibrosis with or without cardiomyocyte hypertrophy. Therefore, the Foz HFD model as well as the long-term high fat high fructose model could be used to explore experimentally the cross talk between the liver and the heart.

We actively explored potential functional impact of the cardiac phenotype. Increased cardiac workload elevates stress on the left ventricular wall, thereby inducing compensatory cardiomyocyte enlargement to support cardiac function. Cardiac hypertrophy triggered by a pathological stimulus is usually one of the factors contributing to the development of heart failure.[31] Foz mice with MASH, but not insulin resistant WT mice fed the similar HFD diet, show enlarged cardiomyocytes and a cardiomyopathy phenotype with circumferential LV hypertrophy. This is accompanied by foetal cardiac reprogramming and myocardial fibrosis. The re-expression of foetal *Myh7* with concomitant downregulation of *Myh6* is a well-known characteristic of pathological cardiac hypertrophy in rodents.[32] Changes in natriuretic peptides expression (ANP) and secretion (BNP), as seen in Foz mice with MASH, also support adverse remodelling and altered myocardial metabolism.[33] In particular the observed elevated levels of plasma BNP are of importance in this context since they are considered as gold standard for both the diagnosis and prognosis of heart failure,[34] into which pathological alterations of the ventricular structure can progress.

In hypertrophied hearts, *Slc2al* expression and basal glucose uptake are enhanced.[35] The same upregulation of *Slc2al* was seen in the hearts of FH mice, supporting insulin resistance, and increased unregulated glucose uptake in the face of permanent hyperglycaemia in the remodelled myocardium of Foz mice with MASH. The observed myocardial fibrosis further supports the hypothesis of pathological remodelling. lmportantly, a similar pattern of ill-adaptive (pathological/unphysiological) cardiac remodelling is recapitulated in a second model of severe MASH, the C57BL/6J mice with a fat- and fructose rich dietary regimen.

Despite the aforementioned alterations of the myocardium, ejection fraction is preserved as documented in Foz mice. Thus, remodelling and fibrosis had no influence on ejection fraction or on fractional shortening, thereby indicating that the contractile function of the myocardium is still intact. lmportantly, heart failure with preserved ejection fraction (HFpEF) is a form of heart failure primarily associated with diastolic dysfunction, frequently encountered in patients with metabolic deregulations. Some phenotypes of HFpEF are even considered as cardiac manifestations of MASLD.[36] Unfortunately, morbid obesity in our animals precluded the acquisition of valid ultrasound data pertaining to diastolic function or atrial morphology. Although LV remodelling and fibrosis in Foz mice with MASH are likely to increase ventricular wall stiffness and consequently decrease myocardial relaxation, analysis utilizing PV loops confirmed that the cardiac function is still undisturbed. Yet, the cardiomyopathy could set the high fat diet fed Foz mice at risk of decompensation. Although we observed no significant aggravation of the ventricular hypertrophy between 12 and 24 weeks, decompensation and failure may come with time. lt was therefore of high interest to verify whether diastolic dysfunction and heart failure might be evident after a longer exposure to high fat diet and liver disease and to examine if aging aggravates the MASH-associated cardiomyopathy. We therefore set up a 60 weeks feeding experiment. By this time, wild type mice had developed signs of aging with cardiac fibrosis. The phenotype was not significantly aggravated by HFD feeding in Foz counterparts. First, we noted that, in aged HFD-fed Foz mice, insulin resistance was alleviated as high insulin production successfully controlled blood glycose homeostasis in aged HFD-fed Foz unlike in younger ones, as shown here in the fasting state. A better glycaemic control may prevent excessive cardiomyocyte glucose uptake and cardiotoxicity.[37] Second, the characteristics of liver disease have changed: rather than steatosis, hepatocyte ballooning and lobular inflammation that characterize MASH in the 24weeks feeding experiment, liver histology after long term HFD in Foz demonstrates the absence of significant steatosis as well as the absence of ballooning although there is considerably more collagen deposition and even hepatocellular carcinoma in some animals. lf the increased CV risk associated with MASH is driven by liver lipotoxicity, then burn-out MASH may rather be more cardio-protective. Whether either or both glycemia and liver disease contribute to the adverse cardiac remodelling we observed, requires further investigation. This could help answering the question if liver-directed therapeutic control of MASH, as already demonstrated for glycaemic control,[38] could reduce CV risk and events.

The cardiomyopathy may predispose Foz mice with MASH, a condition present as early as after 8 weeks of HFD,[39] to heart failure. This proposition is supported by our data. As expected, administration of angiotensin ll triggers an elevation of blood pressure in both WT and Foz mice. Considering that the utilized dosage is the same and that relative heart weight and cardiomyocyte size as well as the extent of the induced hypertension in Foz and WT mice are comparable, too, we can also expect that the intensity of these direct and indirect stressors is equal in the hearts of both groups. However, the observed stronger dysregulation of cardiac remodelling-related gene expression in Foz mice as a result of the Ang ll treatment indicates involvement of an additional effector. In MASLD patients the CVD risk is known to be higher when they suffer from MASH,[40] hence the inflamed liver represents a highly plausible contributor to the stronger cardiac phenotype seen in Foz mice. This interpretation is further supported by the observed strong tendency of left atrial enlargement in mice with MASH, a hallmark of atrial structural remodelling triggered by chronic cardiac volume and pressure overload.[41] In contrast to mice with MASH, those with simple fatty liver show absolutely no sign of atrial remodelling, despite the treatment with Ang ll. Again, this indicates that beside Ang ll a further deleterious factor affects the heart in the presence of MASH. Another affirming argument for our interpretation is the observed reduction in end-diastolic volume in combination with the tendency of decreased left ventricular relaxation capacity in Foz mice; these adverse alterations of cardiac functionality suggest that Foz mice with MASH but not their WT littermates with MAFLD are prone to develop left ventricular diastolic dysfunction.

We are well aware that sole usage of male animals represents a limitation of this study, since both MASLD and CVD feature gender specific differences, with women [19,42] and female mice [43,44] possessing a lower, oestrogen-dependent risk for these diseases compared to their male counterparts. However, we think that the higher risk for both diseases in males, the advantage of excluding a potentially confounding factor in the first cardiac assessment of this model in combination with the circumstance that there is still no established preclinical model for interorgan crosstalk in MASLD and CVD, legitimates this approach. Of course, when transferring future findings obtained with this model to clinical studies, the potential effects of gender differences should then be taken into account.

In conclusion, male mice with MASH, in two unrelated models, also develop a hypertrophic cardiomyopathy with adverse remodelling in the presence of steatohepatitis. However, these clearly pathological changes in the heart do not impact cardiac function in resting mice. As supported by observations in mice challenged with angiotensin ll, increased cardiovascular stress or workload may aggravate myocardial remodelling, and promote heart failure. Our data are also compatible with the proposition that liver lipotoxicity (manifested by steatosis and ballooning) together with heart glucotoxicity (consequence of hyperglycaemia and unregulated increased glucose uptake by the cardiomyocytes) leads to the MASH-associated cardiomyopathy.

## Clinical Perspectives

-The objective of the study was to examine whether metabolic dysfunction-associated steatotic liver disease (MASLD)/metabolic dysfunction-associated steatohepatitis (MASH) promotes cardiomyopathy and cardiac dysfunction, and to assess the suitability of a single pre-clinical model to examine CVD and NAFLD concomitantly

-Mice with MASH, in 2 independent models, develop an adverse cardiac remodelling characterized by a robust transcriptomic signature and cardiac fibrosis, indicating that MASH might actively contribute to the development of pathological structural changes in the heart.

-Therapeutic control on MASH together with improved glucose homeostasis may confer cardioprotection, a hypothesis that is worth testing, given the potential benefits - the two MASH models examined in this study may be of use to explore the liver-heart axis.

## Disclaimer

The authors declare that there are no conflicts of interest.

## Data Availability Statement

Original data are available upon reasonable request.

## Acknowledgements

We thank Caroline Bouzin (2lP/lREC/UCLouvain, Brussels, Belgium) for her expert assistance concerning image analysis. We also thank Natacha Feza-Bingi, Corinne Picalausa and Simon Ravau (GAEN/lREC/UCLouvain, Brussels, Belgium) as well as Aurelie Daumerie (2lP/UREC/ UCLouvain, Brussels, Belgium) for technical support.

## Financial support

This work was supported by funding from the Fonds de la Recherche Scientifique (FNRS) with the project PDR T.0141.19 - SARCONASH to lL and PDR THEMA-CARDlO: “CVD in NASH” P.C006.22 to lL and CD. CD is a senior research associate with the FNRS, Belgium.

## Contributions

SB and JL contributed equally to the work, CD and lL share senior authorship. SB, JL, AM (cardiac catheterization), EPD and HE (both cardiac ultrasound) performed experiments and acquired data. SB, JL, AM, EPD, CB, CD and lL analysed data. CD and lL designed the study, supervised the experiments, and obtained funding. SB and lL drafted the manuscript. All authors reviewed and validated the manuscript.

---

**Supplements, figure 1:**
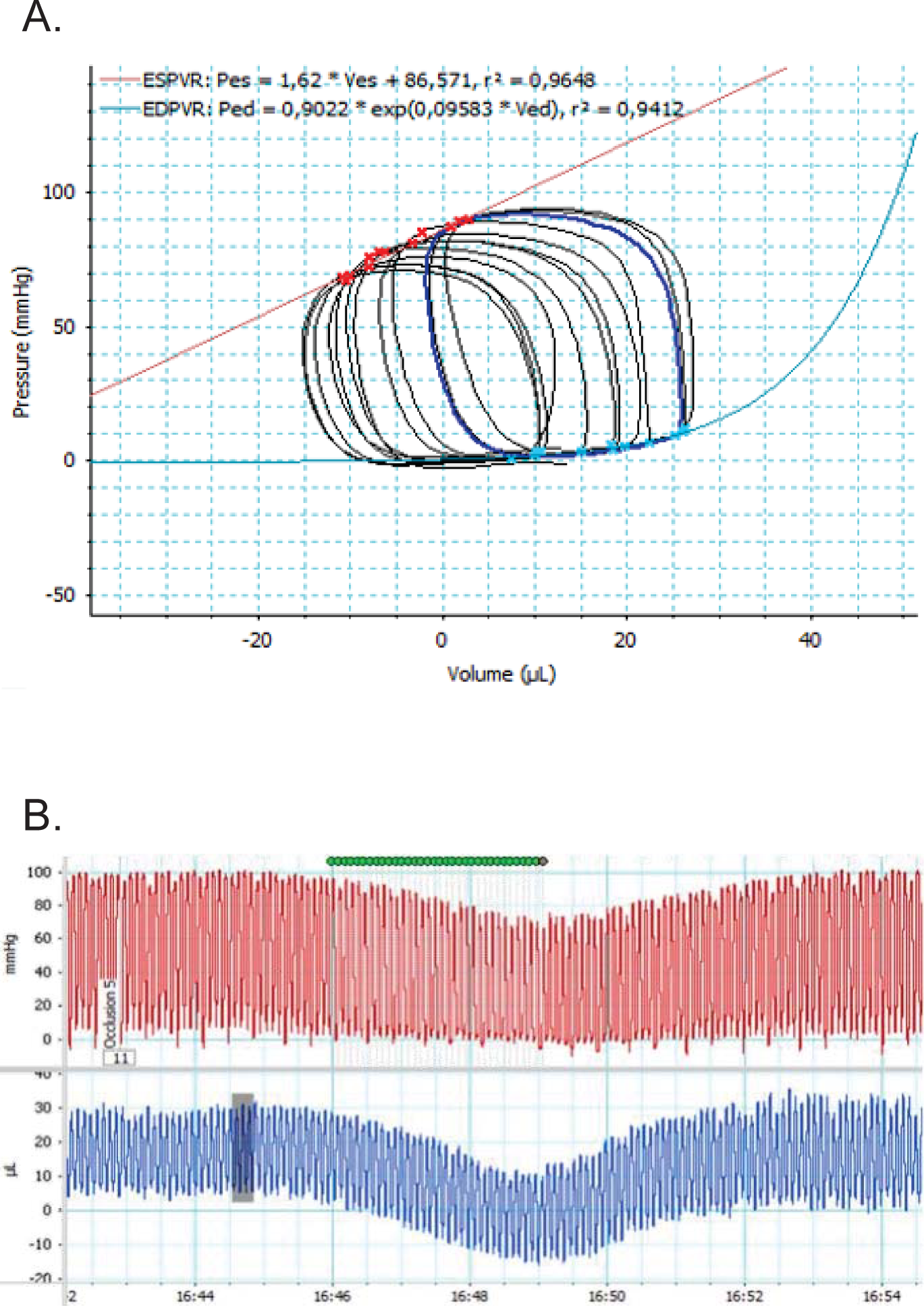
representative PV loop data recording. (A) PV loops during vena cava occlusion in WN mouse; (B) left ventricular pressure (upper part, red) and volume (lower part, blue) changes.

**Supplements, table 1:** in vivo evaluation of heart function in WT/Foz mice by echocardiography. Cardiac parameters were examined after 12 (white fields) and 20 (black fields) weeks of feeding. E/A- and E/e’-ratio of FH mice are not measurable (n.m.) in older animals since exaggerated obesity prevented the acquisition of data of adequate validity. FS was analyzed via M-mode, all remaining parameters were measured using B-mode. **Statistics: n=5-6 per group; the focus of this examination was the potential impact of MASH, thus p-values are displayed for comparison of FH (MASH liver) with WN (healthy liver) values; n.s.=not significant; two-way ANOVA (2wA) with Tukey’s multiple comparisons test; Kruskal-Wallis (K-W) with Dunn’s multiple comparisons test.**

| parameter | WN | WH | FN | FH | p-value | statistical test |
| --- | --- | --- | --- | --- | --- | --- |
| Body weight (BW), g | 26.85 ± 0.54 | 31.15 ± 0.54 | 30.53 ± 1.31 | 55.12 ± 1.22 | *** | K-W |
|  | 28.58 ± 0.93 | 36.07 ± 0.87 | 38.44 ± 2.56 | 63.78 ± 1.49 | *** | K-W |
| End-systolic volume (ESV), µL | 13.70 ± 1.22 | 13.52 ± 1.82 | 11.95 ± 0.73 | 13.66 ± 1.10 | n.s. | K-W |
|  | 12.29 ± 1.60 | 13.51 ± 1.20 | 15.21 ± 1.33 | 12.92 ± 1.18 | n.s. | K-W |
| End-diastolic volume (EDV), µL | 39.23 ± 2.21 | 40.66 ± 2.15 | 39.35 ± 2.26 | 43.69 ± 2.38 | n.s. | K-W |
|  | 40.92 ± 3.95 | 49.40 ± 2.87 | 45.30 ± 2.99 | 49.35 ± 2.01 | n.s. | K-W |
| Ejection fraction (EF), % | 65.10 ± 2.16 | 67.39 ± 3.59 | 69.47 ± 1.39 | 68.76 ± 1.63 | n.s. | K-W |
|  | 70.01 ± 2.13 | 72.83 ± 1.63 | 66.44 ± 1.44 | 73.91 ± 2.09 | n.s. | K-W |
| Fractional shortening (FS), % | 44.67 ± 3.54 | 45.43 ± 2.05 | 44.09 ± 2.23 | 46.15 ± 2.30 | n.s. | K-W |
|  | 44.17 ± 2.70 | 46.44 ± 2.29 | 46.77 ± 2.71 | 49.60 ± 1.52 | n.s. | K-W |
| Left ventricular (LV) mass, mg | 137.60 ± 1.06 | 145.7 ± 9.55 | 144.90 ± 5.11 | 164.2 ± 9.06 | n.s. | 2wA |
|  | 135.30 ± 11.03 | 154.30 ± 10.74 | 154.50 ± 15.65 | 165.60 ± 6.72 | n.s. | 2wA |
| Heart rate (HR), bpm | 566.80 ± 11.20 | 548.60 ± 14.46 | 533.50 ± 14.50 | 526.70 ± 20.22 | n.s. | 2wA |
|  | 552.10 ± 12.27 | 540.30 ± 11.87 | 548.30 ± 13.45 | 469.10 ± 34.53 | * | 2wA |
| Left ventricular posterior wall at end systole (LVPWs), mm | 1.32 ± 0.06 | 1.39 ± 0.07 | 1.35 ± 0.06 | 1.62 ± 0.07 | n.s. | K-W |
|  | 1.37 ± 0.07 | 1.43 ± 0.06 | 1.50 ± 0.06 | 1.68 ± 0.05 | * | K-W |
| Left ventricular posterior wall at end diastole (LVPWd), mm | 0.77 ± 0.03 | 0.82 ± 0.04 | 0.83 ± 0.06 | 0.99 ± 0.05 | * | K-W |
|  | 0.81 ± 0.03 | 0.83 ± 0.03 | 0.88 ± 0.03 | 0.98 ± 0.04 | * | K-W |
| Interventricular septum at end systole (IVSs), mm | 1.23 ± 0.02 | 1.27 ± 0.06 | 1.27 ± 0.06 | 1.49 ± 0.07 | * | 2wA |
|  | 1.24 ± 0.06 | 1.40 ± 0.08 | 1.43 ± 0.08 | 1.59 ± 0.06 | * | 2wA |
| Interventricular septum at end diastole (IVSd), mm | 0.82 ± 0.02 | 0.87 ± 0.04 | 0.84 ± 0.05 | 1.00 ± 0.06 | * | 2wA |
|  | 0.84 ± 0.05 | 0.93 ± 0.05 | 0.91 ± 0.05 | 1.07 ± 0.08 | * | 2wA |
| Left ventricular internal diameter at end-systole (LVIDs), mm | 1.86 ± 0.16 | 1.86 ± 0.10 | 1.92 ± 0.12 | 1.95 ± 0.12 | n.s. | 2wA |
|  | 1.87 ± 0.11 | 1.97 ± 0.12 | 1.93 ± 0.10 | 1.93 ± 0.08 | n.s. | 2wA |
| Left ventricular internal diameter at end-diastole (LVIDd), mm | 3.33 ± 0.08 | 3.39 ± 0.08 | 3.42 ± 0.11 | 3.63 ± 0.14 | n.s. | 2wA |
|  | 3.34 ± 0.07 | 3.67 ± 0.10 | 3.64 ± 0.10 | 3.82 ± 0.10 | ** | 2wA |
| E/A ratio | 2.16 ± 0.26 | 1.82 ± 0.15 | 2.01 ± 0.13 | 1.69 ± 0.23 | n.s. | 2wA |
|  | 2.55 ± 0.23 | 2.09 ± 0.31 | 2.84 ± 0.33 | n.m. | - | - |
| E/e' ratio | 32.33 ± 0.54 | 35.19 ± 2.66 | 27.36 ± 0.61 | 26.47 ± 6.53 | n.s. | K-W |
|  | 29.97 ± 1.39 | 31.37 ± 1.82 | 29.09 ± 3.12 | n.m. | - | - |

## Notes

### Competing Interest Statement

The authors have declared no competing interest.

